# DeepProfile: Deep learning of cancer molecular profiles for precision medicine

**DOI:** 10.1101/278739

**Authors:** Ayse Berceste Dincer, Safiye Celik, Naozumi Hiranuma, Su-In Lee

## Abstract

We present the DeepProfile framework, which learns a variational autoencoder (VAE) network from thousands of publicly available gene expression samples and uses this network to encode a low-dimensional representation (LDR) to predict complex disease phenotypes. To our knowledge, DeepProfile is the first attempt to use deep learning to extract a feature representation from a vast quantity of unlabeled (i.e, lacking phenotype information) expression samples that are not incorporated into the prediction problem. We use DeepProfile to predict acute myeloid leukemia patients’ *in vitro* responses to 160 chemotherapy drugs. We show that, when compared to the original features (i.e., expression levels) and LDRs from two commonly used dimensionality reduction methods, DeepProfile: (1) better predicts complex phenotypes, (2) better captures known functional gene groups, and (3) better reconstructs the input data. We show that DeepProfile is generalizable to other diseases and phenotypes by using it to predict ovarian cancer patients’ tumor invasion patterns and breast cancer patients’ disease subtypes.

## 1. Introduction

Learning robust prediction models based on molecular profiles (e.g., expression data) and complex phenotype data (e.g., drug response) is a crucial step toward realizing the many benefits of personalized medicine. However, most expression datasets are high-dimensional (i.e., #samples ≪ #variables) and therefore, it is challenging to use them to learn accurate prediction models. Learning a function that maps observed molecular features to an informative lowdimensional representation (LDR) is the key to success in overcoming the bane of dimensionality.

We present DeepProfile, which uses VAEs to learn an unsupervised neural network model of gene expression from thousands of cancer patients, and then uses this model to encode an LDR to predict complex phenotypes of patients excluded from network training. To our knowledge, DeepProfile is the first attempt to predict complex phenotypes using an LDR based on an unsupervised deep learning-based method that is completely blind to both expression and phenotype data from the test samples. DeepProfile has three unique aspects. First, since DeepProfile learns an unsupervised model in the training step, it can use a vast quantity of samples from which only gene expression data is available. Second, it can encode an LDR for a new cancer patient by transferring network information learned by the trained model from a much greater number of individuals. Finally, this newly encoded LDR can be effectively used to predict any phenotype information for the new patient.

Since it learns an LDR based on a deep neural network (DNN), DeepProfile has a potential to capture complex, non-linear relationships between expression and phenotype data. Consistent with that, our experimental results show that DeepProfile better reveals hidden structures within the data. It highly improves prediction of *in vitro* drug response of acute myeloid leukemia (AML) patients to 160 chemotherapy drugs when compared to the original features (i.e., expression levels) and LDRs learned based on two commonly used dimensionality reduction methods – PCA and k-means clustering – trained on the same large set of samples DeepProfile is trained on. Moreover, the LDR learned based on DeepProfile can reconstruct input expression data and capture known gene pathways more accurately than LDRs based on PCA or k-means. Consistent results from additional applications of DeepProfile to ovarian cancer (OV) to predict tumor histopathology and breast cancer (BRCA) to predict patient subtypes imply that DeepProfile is a general framework applicable to any cancer with a large number of publicly available samples and various cancer phenotypes.

## 2. Related Work

We first describe studies that applied unsupervised deep learning to expression data, then we mention machine learning methods used for drug response prediction, the main task which led us to design and build the DeepProfile framework.

Tan et al. (2014) used denoising autoencoders and examined the learned LDR in terms of its association with known breast cancer features. Others used different autoencoder models to learn an LDR for gene expression to improve the clustering performance (Gupta et al., 2015; Cui et al., 2017).

Rampasek et al. (2017) built semi-supervised VAE models to improve drug response prediction accuracy using pre- and post-treatment cell lines. Way & Greene (2017) used VAE to learn biologically relevant latent space from The Cancer Genome Atlas (TCGA) pan-cancer data. Chiu et al. (2018) used autoencoders to predict drug response for various cancers using expression and mutation data. DeepProfile is different from all these approaches because it is the first to predict complex cancer phenotypes using an LDR learned based on a VAE trained from almost all of the available GEO patient expression samples for a cancer.

Drug response prediction has been addressed by several authors most commonly by using ridge or elastic net regression (Garnett et al., 2012; Barretina et al., 2012; Jang et al., 2013). Several other studies used more complex models such as support vector machine and random forest to improve prediction accuracy (Stetson et al., 2014) as well as multitask learning (Costello et al., 2014; Yuan et al., 2016). DeepProfile is separated out from these studies as it transfers information from many patients with the same cancer type using a deep autoencoder to predict drug response. Also, DeepProfile uses only a small subset of genes while past studies show performance on the entire set of genes.

## 3. Methods

### 3.1. The DeepProfile Framework

As shown in Fig. 1, DeepProfile: (1) learns a network representation from the gene expression of thousands of AML patients in an unsupervised way, (2) uses the learned network to encode an LDR for 30 held-out AML patient samples whose *in vitro* responses to 160 drugs are available, and (3) predicts the drug response of these patients using the encoded LDR. DeepProfile adopts a VAE (Kingma & Welling, 2013) to learn a network representation for the gene expression. VAE, an extension of a standard autoencoder (AE), is an unsupervised DNN which uses variational inference to infer the posterior distribution of latent embeddings. A VAE’s objective is to minimize the error between the input data and the data reconstructed from the embeddings; but it also assumes that the posterior is normally distributed. It adopts a standard Gaussian prior on the embeddings, which generally yields an LDR more relevant and informative about the input data, and more generalizable to unseen data compared to an LDR learned by an AE. Our VAE model consists of encoder and decoder networks, each with three dense layers. We used sum of mean squared error (MSE) between input and reconstructed data and Kullback-Leibler (KL) divergence between the posterior and prior as an objective function and trained using Adam method (Kingma & Ba, 2014). We built our VAE model, available at https://github.com/suinleelab/DeepProfile, using Keras with Tensorflow backend.

**Figure 1.**
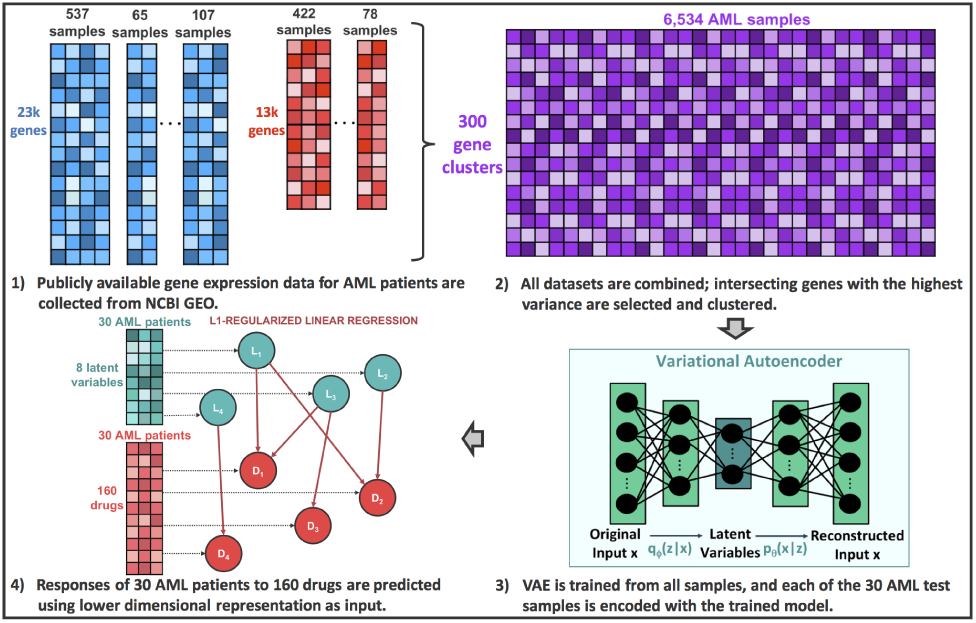
The DeepProfile framework. We combined standardized expression data collected from 96 studies and corrected batch effects both within each study and across studies. The genes with a median absolute deviation (MAD) higher than the mean MAD are selected and clustered. The VAE network is trained using a total of 300 gene clusters of 6,534 samples. Using the trained network, an 8-dimensional LDR is learned for each of the 30 AML patient samples, for which we have the *in vitro* response to 160 drugs. We use the LDR as an input of an L1-regularized linear regression to predict the response to each drug.

### 3.2. Data Collection and Experimental Setup

Genome-wide gene expression data consists of thousands of variables (i.e., genes) and requires a large number of training samples to learn a network representation that is relevant to the biological mechanisms and generalizable to broader populations. Thus, we trained our VAE models using a large number of publicly available expression samples that we collected from the National Center for Biotechnology Information (NCBI) Gene Expression Omnibus (GEO) database (Edgar et al., 2002) using a script we coded to automatically download GEO data. We used microarray data from two platforms with the highest number of samples in GEO – Affymetrix HG-U133 plus 2.0 and Affymetrix HG-U133A. Excluding healthy samples and cell lines resulted in a total of 6,534 patient samples to be used by DeepProfile. To our knowledge, we included all available AML patient samples in GEO from the aforementioned two platforms.

To integrate data from various platforms, we took 13,237 genes available for all datasets. A study might have different sample batches submitted on different dates. Therefore, we corrected for the potential batch effects within each study using ComBat (Johnson et al., 2007). We then standardized (i.e., made zero-mean and unit variance) each gene in each dataset to ensure that different input features (here, gene expression levels) were on the same scale. We applied batch effect correction to the entire data set once again using ComBat, considering each study to be a separate batch in order to minimize the effect of potential confounders due to experimental variations. We then removed the genes that had a median absolute deviation (MAD) below the mean MAD. As a result, 5,393 genes were selected, which we divided into 300 clusters (using agglomerative clustering) so that those with similar expression patterns were grouped together, reducing the noise and dimensionality of the feature space. We used centroids of the learned 300 clusters to train the VAE model. Then we used the network learned by VAE to encode an 8-dimensional LDR for each of the 30 AML patient samples from Lee and Celik et al. (2018), which were collected by the University of Washington Medical Center and measured in terms of genome-wide gene expression and *in vitro* response to 160 chemotherapy drugs. We used the encoded LDR in an L1-regularized linear regression setting and measured drug response prediction performance for these patients separately for each drug. We computed the prediction error using a cross-validation (CV) test and also performed an additional CV on training samples to select the regularization parameter *λ*. Since the VAE model is non-convex, the learned LDR is not unique and may depend on the initial network weights. To ensure that our results considered the potential variation in the prediction performance due to the variation in the learned LDRs, we trained the VAE model ten times and retrained the prediction models for each of the ten different 8-dimensional LDRs. Our result figures include the error bars representing one standard deviation across predictions.

Next, we applied DeepProfile to additional two cancers and used the learned LDRs to predict tumor invasion pattern for OV and tumor subype for BRCA. For each of these additional applications, we applied the exact same procedures for GEO data collection and preprocessing and CV tests as for AML application. To our knowledge, for these cancers as well, we used all GEO samples from the same two platforms we used for AML. The only difference is that we used a logistic regression since these are classification tasks.

#### Additional applications - OV and BRCA

We trained VAE models for OV using a total of 2,714 training samples and 300 clusters learned from 4,572 genes with a MAD higher than the mean MAD. Then, using the trained network, we encoded an LDR for the expression of 85 TCGA OV samples (Network, 2011) for which we have the invasion pattern phenotype (infiltrative or expansile) that was provided by Celik et al. (2016). We then predicted the invasion pattern phenotype using the encoded LDR. Similarly, we trained VAE models for BRCA using a total of 11,963 training samples and 300 clusters learned from 3,048 genes with a MAD higher than the mean MAD. Then, using the trained network, we encoded an LDR for the expression of 493 TCGA BRCA samples and predicted these samples’ subtypes (Basal-like, Her2-enriched, Luminal-A, Luminal-B) provided by TCGA (Network, 2012).

## 4. Results

We compared DeepProfile to the original gene expression features and LDRs from two commonly used dimensionality reduction methods – PCA and k-means – applied to the same set of 6,534 AML patient samples from GEO. We evaluated our method in three ways: (1) measuring how well the learned LDR predicts drug response, (2) analyzing how learned latent variables match known, biologically meaninful gene pathways, and (3) computing the Spearman correlation between the original and VAE-reconstructed data for each sample in the training and test sets.

We first compared the MSE obtained by DeepProfile trained from all microarray samples to the gene expression levels of 30 AML test patients and 8 LDR features learned by PCA and k-means. The goal of this experiment is to determine whether the transferred network learned in an unsupervised manner from large amounts of data from a cancer type would help with the task of phenotype prediction for new patients with the same type of cancer. For k-means clustering, we learned 8 gene clusters and used the cluster centroids as LDR, while for PCA, we used top 8 principal components. Fig. 2a compares MSEs averaged over all 160 drugs available in the dataset, and MSEs averaged over the subset of drugs commonly used for AML treatment as provided by Lee and Celik et al. (2018); DeepProfile outperformed all other methods for both set of drugs.

**Figure 2.**
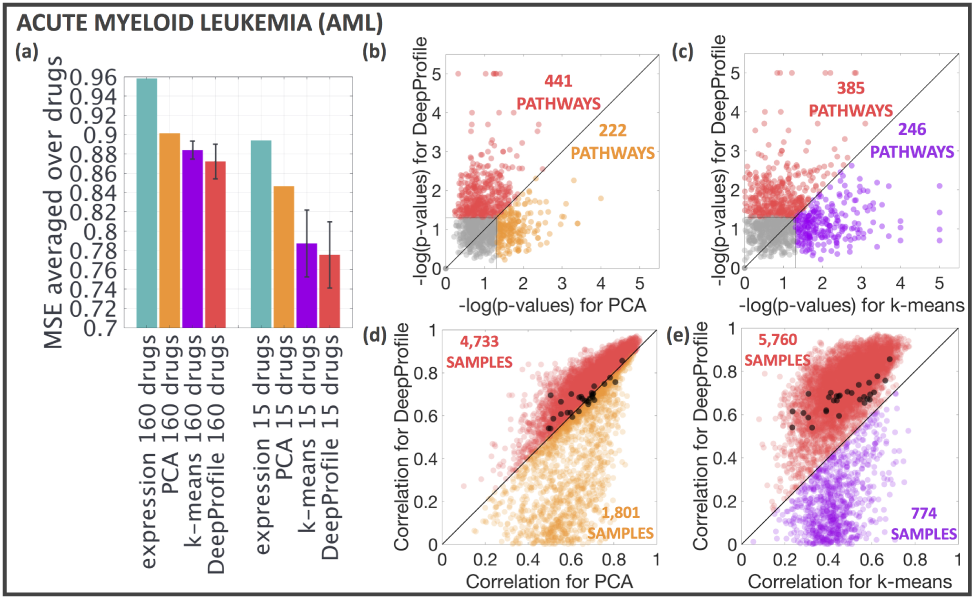
(a) Comparison of drug response prediction MSEs averaged over all 160 drugs or 15 drugs commonly used for AML patients. (b), (c) Comparison of pathway recovery performance between DeepProfile and (b) PCA and (c) k-means. Each dot represents one of the 1,077 pathways. Horizontal and vertical lines represent the statistical significance threshold *p* = 0.05. (d), (e) Scatter plot of Spearman correlation between the original and reconstructed expression comparing DeepProfile to (d) PCA and (e) k-means. Test samples are marked in black.

Next, we checked whether the genes that belong to known functional pathways (1,077 Reactome, BioCarta, and KEGG GeneSets from the C2 collection of the current version of MSigDB (Subramanian et al., 2005) are highly ranked by DeepProfile. We computed the weights of each gene for each of the 8 LDR features (i.e., how much each gene contributed to the value of each LDR feature) learned based on PCA, k-means, or Deep-Profile. We determined gene weights using the Keras implementation of the Integrated Gradients method (Sundarara-jan et al., 2017) provided at https://github.com/hiranumn/IntegratedGradients. We computed the ranking of each gene based on its weight magnitude and performed a permutation test to check whether the average ranking of the genes in the pathway was higher than it would be by a random chance. We performed 10,000 random permutations for each of the methods, and compared DeepProfile to PCA and k-means in terms of the permutation p-values. Of the well-captured pathways (i.e., the ones that are captured by at least one of the methods with a p-value *<* 0.05), 441 were better captured by DeepProfile than by PCA (Fig. 2b) and 385 were better captured than by k-means (Fig. 2c). For each pathway, we performed the permutation test for the top LDR feature from each method (i.e., the embedding with the highest average ranking of the pathway genes) because it is common that in a DNN model like VAE, each hidden node captures a separate functional unit that contributes to the learned meaningful representation of the data. Thus, we believe that it is reasonable to assume that each pathway, which can be viewed as a functional unit of gene expression, is represented by the LDR feature that leads to the highest average ranking of the genes in it.

Fig. 2d, e compare the Spearman correlation between the original and reconstructed AML training and test samples. DeepProfile achieved a better reconstruction correlation for both training and test samples than PCA or k-means. Since k-means provides a hard assignment of genes to clusters, we could not use any cluster membership weights of the genes while reconstructing data from the cluster centroids. That is likely the reason for the significantly lower reconstruction performance observed for k-means than for PCA.

### Results from OV and BRCA

When used for OV tumor invasion classification (Fig. 3a) and BRCA tumor subtype classification (Fig. 3f), DeepProfile again achieved a lower classification error than the original expression features of the test samples and LDRs based on PCA and k-means trained on the same set of samples as DeepProfile. DeepProfile trained for OV could capture 372 pathways significantly better than PCA (Fig. 3b) and 353 pathways significantly better than k-means (Fig. 3c). It also outperformed these two methods in terms of the data reconstruction performance (Fig. 3d, e). Similarly, DeepProfile trained for BRCA could capture 314 pathways significantly better than PCA (Fig. 3g) and 364 pathways significantly better than k-means (Fig. 3h), and again outperformed these two methods in terms of the data reconstruction performance (Fig. 3i, j).

**Figure 3.**
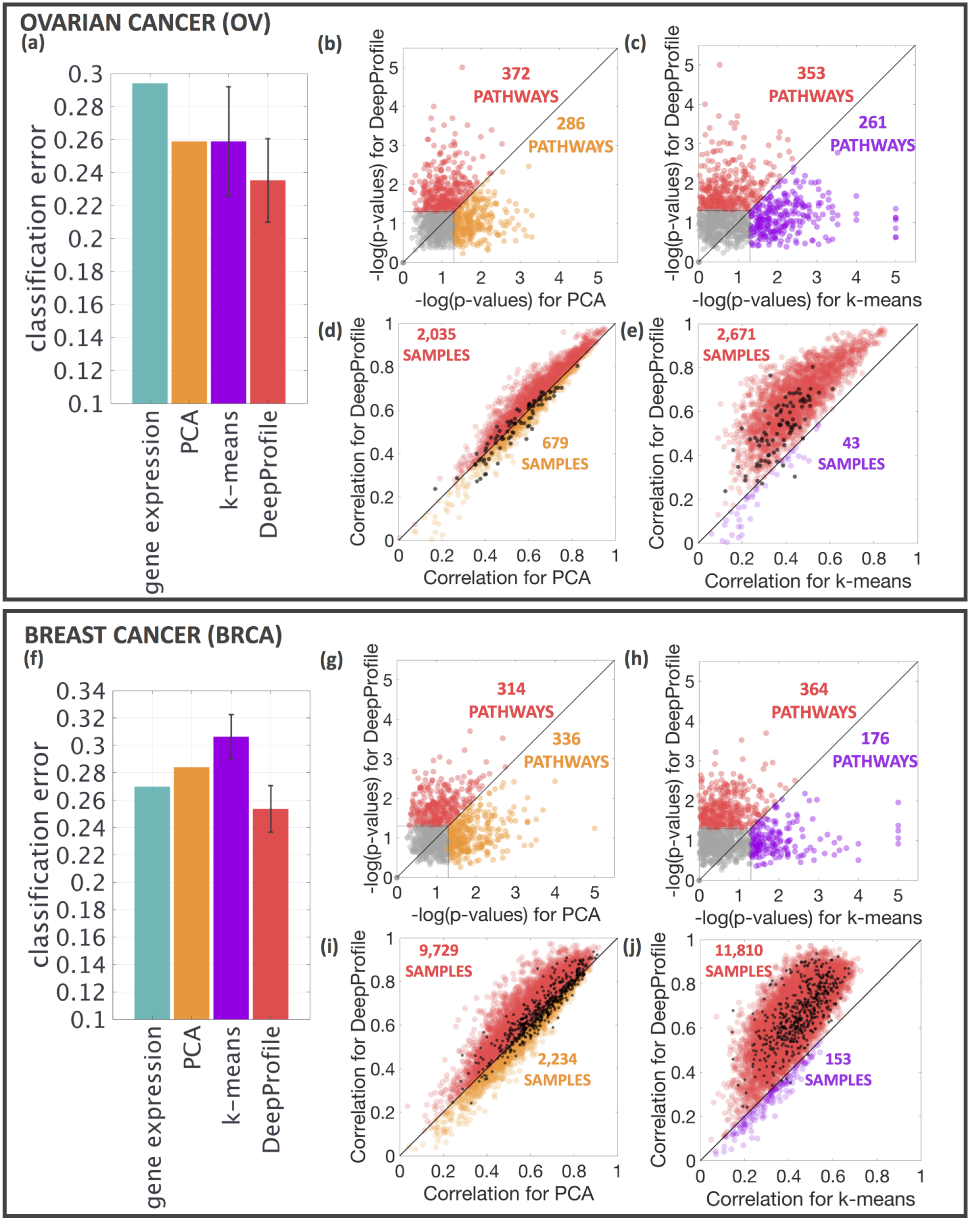
OV and BRCA results. (a) and (f) compare the error obtained by different methods for classifying the OV tumor invasion patterns and BRCA tumor subtypes, respectively. See the text or Fig. 2 caption for the details of the scatter plots in (b-e) and (g-j).

## 5. Discussion

We present the DeepProfile framework, which adopts a variational autoencoder (VAE) to learn a network representation (LDR) from publicly available, unlabeled gene expression data and uses that to predict drug response of new AML patients. Although expression samples used in VAE training were obtained from many different studies, DeepProfile successfully disentangles data discrepancies and learns an informative LDR that accurately predicts complex phenotypes for different cancers.

Our future directions include but are not limited to: (1) training DeepProfile using samples from different cancers which would significantly increase the sample size, thus, statistical power, and yield a resulting model which could be used for predicting phenotypes from different cancers, and (2) extending the scope of DeepProfile by using multi-omics data to create more informative embeddings for cancer.

## References

Barretina, J., Caponigro, G., Stransky, N., Venkatesan, K., Margolin, A. A., Kim, S.,…, and Garraway, L. A. The cancer cell line encyclopedia enables predictive modelling of anticancer drug sensitivity. Nature, 492(7428): 603–7, 2012.

Celik, S., Logsdon, B. A., Battle, S., Drescher, C. W., Rendi, M., Hawkins, R. D., and Lee, S. Extracting a low-dimensional description of multiple gene expression datasets reveals a potential driver for tumor-associated stroma in ovarian cancer. Genome Medicine, 8(1):66, 2016.

Chiu, Y., Chen, H. H., Zhang, T., Zhang, S., Gorthi, A., Wang, L., Huang, Y., and Chen, Y. Predicting drug response of tumors from integrated genomic profiles by deep neural networks. arXiv preprint arXiv:1805.07702, 2018.

Costello, J. C., Heiser, L. M., Georgii, E., Gnen, M., Menden, M. P., Wang, N. J.,…, and Stolovitzky, G. A community effort to assess and improve drug sensitivity prediction algorithms. Nature Biotechnology, 32:1202–1212, 2014.

Cui, H., Zhou, C., Dai, X., Liang, Y., Paffenroth, R., and Korkin, D. Boosting gene expression clustering with system-wide biological information: A robust autoen-coder approach. bioRxiv, 2017. doi:10.1101/214122.

Edgar, R., Domrachev, M., and Lash, A. E. Gene expression omnibus: Ncbi gene expression and hybridization array data repository. Nucleic Acids Research, 30(1):207–10, 2002.

Garnett, M. J., Edelman, E. J., Heidorn, S. J., Greenman, C. D., Dastur, A., Lau, K. W.,…, and Benes, C. H. Systematic identification of genomic markers of drug sensitivity in cancer cells. Nature, 483(7391):570–575, 2012.

Gupta, A., Wang, H., and Ganapathiraju, M. Learning structure in gene expression data using deep architectures, with an application to gene clustering. 2015 IEEE International Conference on Bioinformatics and Biomedicine (BIBM), pp. 1328–1335, 2015.

Jang, I. S., Neto, E. C., Guinney, J., Friend, S. H., and Margolin, A. A. Systematic assessment of analytical methods for drug sensitivity prediction from cancer cell line data. In Biocomputing 2014, pp. 63–74, 2013.

Johnson, W. E., Li, C., and Rabinovic, A. Adjusting batch effects in microarray expression data using empirical bayes methods. Biostatistics, 8(1):118–27, 2007.

Kingma, D. P. and Ba, J. Adam: a method for stochastic optimization. arXiv preprint arXiv:1412.6980, 2014.

Kingma, D. P. and Welling, M. Auto-encoding variational bayes. arXiv preprint arXiv:1312.6114, 2013.

Lee, S., Celik, S., Logsdon, B. A., Lundberg, S. M., Martins, T. J., Oehler, V. G.,…, and Becker, P. S. A machine learning approach to integrate big data for precision medicine in acute myeloid leukemia. Nature Communications, 9 (1):42–54, 2018.

Network, The Cancer Genome Atlas. Integrated genomic analyses of ovarian carcinoma. Nature, 474:609–615, 2011.

Network, The Cancer Genome Atlas. Comprehensive molecular portraits of human breast tumours. Nature, 490: 61–70, 2012.

Rampasek, L., Hidru, D., Smirnov, P., Haibe-Kains, B., and Goldenberg, A. Dr.vae: Drug response variational autoencoder. arXiv preprint arXiv:1706.08203, 2017.

Stetson, L. C., Pearl, T., Chen, Y., and Barnholtz-Sloan, J. S. Computational identification of multi-omic correlates of anticancer therapeutic response. BMC Genomics, 15(7): S2, 2014.

Subramanian, A., Tamayo, P., Mootha, V. K., Mukherjee, S., Ebert, B. L., Gillette, M. A.,…, and Mesirov, J. P. Gene set enrichment analysis: A knowledge-based approach for interpreting genome-wide expression profiles. Proceedings of the National Academy of Sciences, 102(43): 15545–15550, 2005.

Sundararajan, M., Taly, A., and Yan, Q. Axiomatic attribution for deep networks. arXiv preprint arXiv:1703.01365, 2017.

Tan, J., Ung, M., Cheng, C., and Greene, C. S. Unsupervised feature construction and knowledge extraction from genome-wide assays of breast cancer with denoising autoencoders. In Biocomputing 2015, pp. 132–143, 2014.

Way, G. P. and Greene, C. S. Extracting a biologically relevant latent space from cancer transcriptomes with variational autoencoders. In Biocomputing 2018, pp. 80–91, 2017.

Yuan, H., Paskov, I., Paskov, H., González, A. J., and Leslie, C. S. Multitask learning improves prediction of cancer drug sensitivity. Scientific reports, 6:31619, 2016.

